# Arylsulfonamide mediated RBM39 degradation causes aberrant splicing of mitotic kinesins

**DOI:** 10.1101/2021.02.01.428819

**Authors:** Seemon Coomar, Alexander Penson, Jürg Schwaller, Omar Abdel-Wahab, Dennis Gillingham

## Abstract

Certain arylsulfonamides (ArSulfs) induce an interaction between the E3 ligase substrate adaptor DCAF15 and the critical splicing factor RBM39, ultimately causing its degradation. Although molecules like the ArSulfs, which interfere with splicing decisions, are exciting potential medicines, the molecular glue mechanism of RBM39 degradation introduces complex pleiotropic effects that are difficult to untangle. For example, DCAF15 inhibition, RBM39 degradation, and the downstream proteome effects of splicing changes will all cause different yet overlaid effects. As such the precise cell-killing mechanism by RBM39 loss is largely unknown. By overlaying transcriptome and proteome datasets, we distinguish transcriptional from post-transcriptional effects, pinpointing those proteins most impacted by splicing changes. Our proteomic profiling of several ArSulfs suggests a selective DCAF15/ArylSulf/RBM39RRM2 interaction with a narrow degradation profile. We identify two mitotic kinesin motor proteins that are aberrantly spliced upon RBM39 degradation, and we demonstrate that these are likely contributors to the antiproliferative activity of ArSulfs.

## Introduction

Recent techniques that use small molecules to co-opt natural mechanisms for regulating protein stability,[1] are exciting innovations because they enable drugs that mimic genetic techniques such as RNAi or CRISPR/Cas9.[2–4] The ubiquitin proteasome pathway (UPP) is one of the primary natural mechanisms for the controlled degradation of proteins.[5] Within the UPP the key molecular signal for protein degradation is the sequential transfer of ubiquitin (Ub) proteins (typically K48-linked Ub 4-6mers) onto client substrates. By binding the surface of an E3 ligase substrate receptor, certain small molecules can act as molecular glues that favor the association and degradation of non-native substrates, while potentially inhibiting the association of natural substrates. Here we study the effects of arylsulfonamide (ArSulf) molecular glues, which degrade the splicing factor RBM39 by directing it to the E3 ligase substrate receptor DCAF15 (**Figure 1A**).

**Figure 1.**
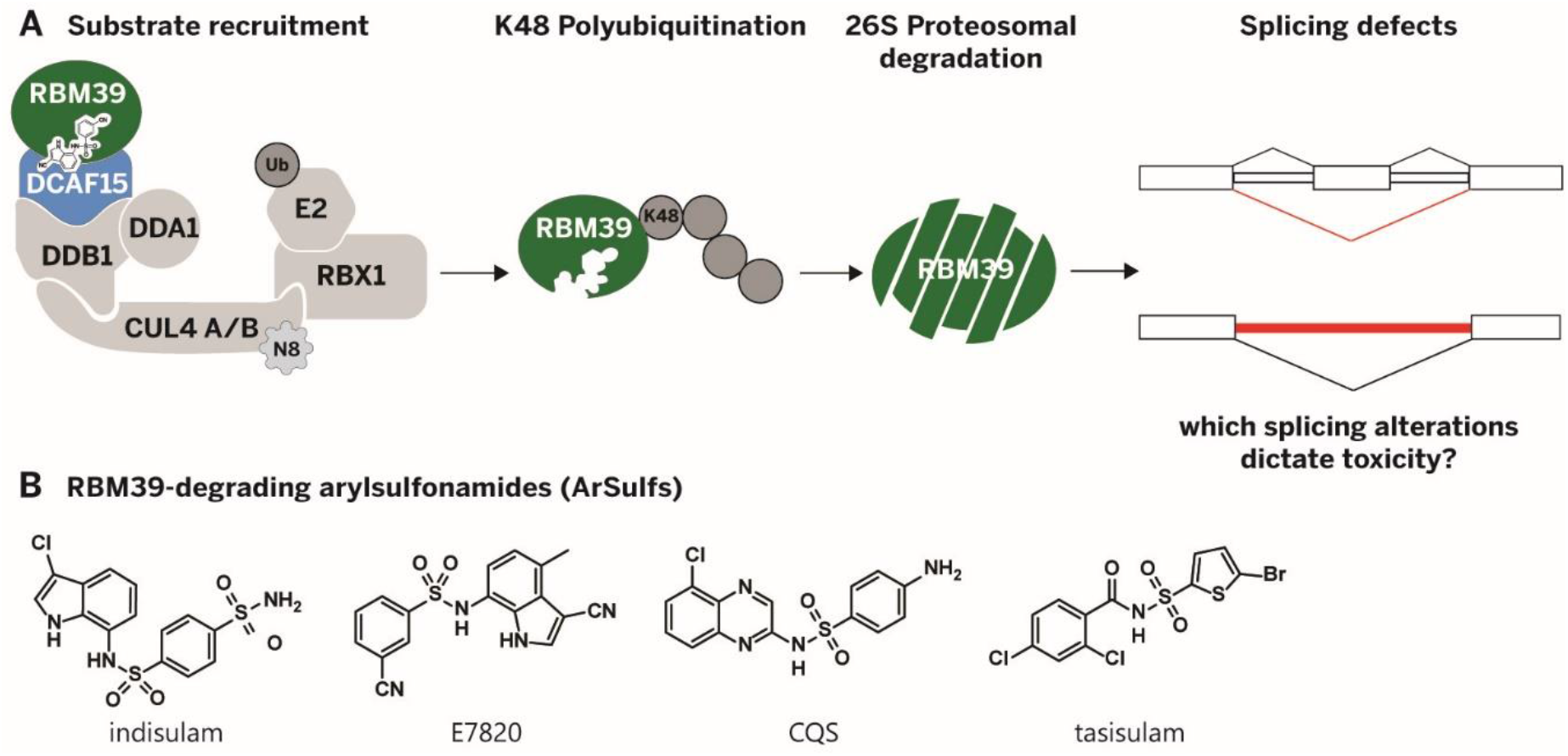
**A)** Arylsulfonamides (ArSulfs) cause the degradation of the splicing factor RBM39 by recruiting it to the DCAF15 cullin-ring ligase; loss of RBM39 causes splicing defects. **B)** The arylsulfonamide compounds proven to degrade RBM39.

Indisulam (**Figure 1B**) is an antitumor arylsulfonamide (ArSulf) that was discovered and tested in clinical trials more than a decade ago,[6–12] but whose mode-of-action was only recently clarified in two independent studies.[13, 14] Several structural studies have since shown that ArSulfs create a new protein-protein interaction by simultaneously binding both DCAF15 and the splicing factor RBM39 (**Figure 1A**).[15–17] Since DCAF15 is an E3 ligase adaptor protein, this binding makes RBM39 susceptible to ubiquitination and subsequent degradation in a mechanism that is reminiscent of the more established immunomodulatory drugs (IMiDs) thalidomide and lenalidomide.[18, 19] The ArSulfs chloroquinoxaline sulfonamide (CQS), tasisulam, and E7820 (**Figure 1B**) are structurally related to indisulam and have all been shown to degrade RBM39. Recent structural and biochemical efforts have identified the second of the two RNA-recognition motifs (RRM) within RBM39 to be responsible for the DCAF15/ArSulf/RBM39 interaction.[15–17, 20, 21] Improving the ArSulf molecular glues could take three forms: enhance DCAF15 binding, increase RBM39 degradation capacity, or expanding the glue interaction to target other RRM-motif bearing proteins. Such improvements are hamstrung by the pleiotropic effects induced in the wake of RBM39 degradation. For example, ArSulfs directly cause DCAF15 inhibition,[22] RBM39 degradation, and downstream transcriptome and proteome effects that result from aberrant splicing. Methods to analyze each of these mechanisms independently will be critical in understanding RBM39 biology, in redeploying DCAF15 against new targets, and in adapting ArSulfs to degrade other RNA binding proteins. We present here a series of experimental and bioinformatics analyses that disentangle the polypharmacological effects of ArSulfs. Our work uncovers RBM39 as a regulator of the mitotic kinesin motor proteins KIF20A and KIF20B, which likely plays a role in the antiproliferative activity of ArSulfs.

## Results

### Assessing proteome-wide selectivity of the DCAF15/ArSulf/RBM39-mediated degradation

To assess the overall proteome effects of the ArSulfs (**Figure 1B**) we performed quantitative TMT proteomics experiments in HCT-116, Mv4;11, and MOLM-13 human cancer cells upon exposure to several ArSulfs. In each cell line the data confirmed the exquisite selectivity of the ArSulfs for RBM39 degradation (**Figure 2A-B, 3A, Figure S1A, S4A**). RBM23 – whose second RRM domain (RBM23_RRM2_) shares near perfect sequence homology with that of RBM39 (RBM39_RRM2_) – as well as RBM5 were the only other RNA binding protein consistently downregulated (**Figure 2A,C**). Although RBM23 is a *bona fide* [17, 21] molecular glue substrate of DCAF15/ArSulfs, it was obscured in the volcano plots by number of other significant hits that were likely not DCAF15 neosubstrates. Since global proteomics analyses incorporate a summation of both transcriptional and post-transcriptional effects, we reasoned that removing RBM39-eCLIP enriched targets (dataset previously generated in one of our labs) [23] from those identified in proteomics might provide a clearer picture of potential molecular glue targets. This strategy enabled an uncluttered look at proteins with lower q-values (q<0.01) that might nevertheless be glue substrates (see filtered volcano plot in **Figure 2B**). Moreover, it removed RBM5, leaving RBM23 as the only other protein containing domains with an RRM (**Figure 2C**). Our results from qPCR measurements in siRNA mediated RBM39 and RBM23 knockdown in HCT-116 cells showed that neither of these was regulating the other on a transcriptional level (**Figure 2D**). Similarly, ArSulf treatment in HCT-116 cells showed no difference in RBM23 transcript levels (**Figure S1C**). For control experiments we generated a DCAF15 double knockout cell line (HCT116^DCAF15−/−^) using CRISPR/Cas9. Treating this cell line with the ArSulfs showed no degradation of RBM23 in TMT-proteomics and confirmed the DCAF15 dependency (**Figure S1B**). In line with recent reports, [15, 17] these results suggest that DCAF15/ArSulf/RBM39 (or RBM23) interaction strictly requires the sequence of the RRM2 domain present in RBM39 and RBM23, confirming ArSulfs as highly selective degraders. Consistent with previous observations [23], we also observed reduced protein levels of MCL1 in the leukemia cell line Mv4;11.

**Figure 2.**
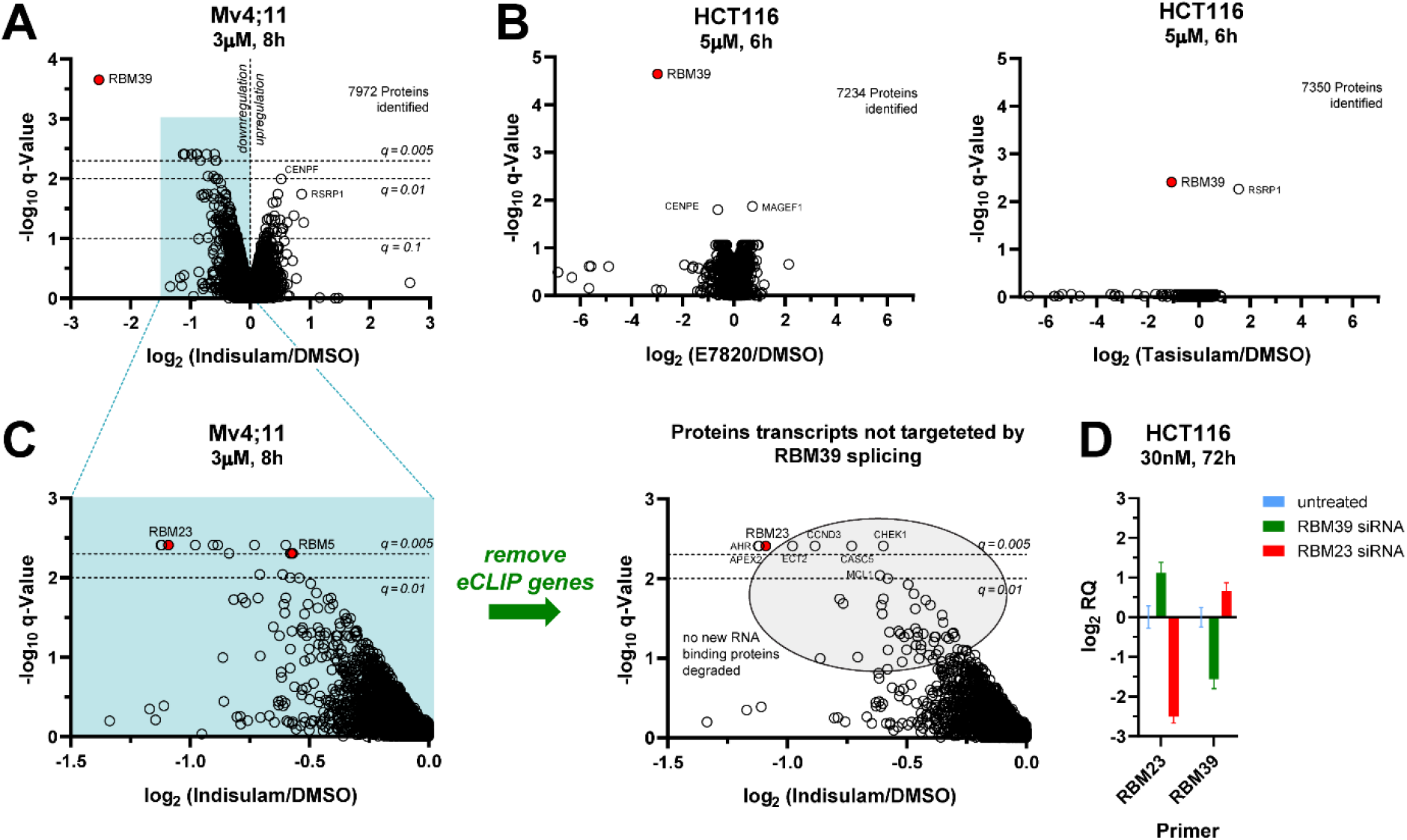
**A)** Change in protein levels relative to vehicle treatment (DMSO) in Mv4;11 cells treated with Indisulam at 3 μM for 8 h quantified by TMT labelling and LC MS/MS analysis, versus q-Value (Benjamini-Hochberg corrected p-Value from Bayes moderated t-statistics). **B)** (left) As for A) but in HCT-116 cells treated with the sulfonamides at 5 μM for 6 h (see also Figure S1A-B). **C)** Proteins retained after removing genes identified in RBM39-eCLIP. **D)** qRT-PCR measurement showing the change in amplicon levels of RBM23 and RBM39 in HCT-116 cells exposed to siRNA mediated RBM39 and RBM23 knockdown (72 h, 30 nM) to untreated HCT-116 cells (conducted in technical triplicates) (see also Figure S1C). The quantification was done by using StepOne Software v2.3 to calculate RQ (same as 2^-ΔΔCt^) relative to DMSO using GAPDH as the housekeeping gene. The error bars were calculated from RQMax and RQMin as defined in the software using 95% CI.

**Figure 3.**
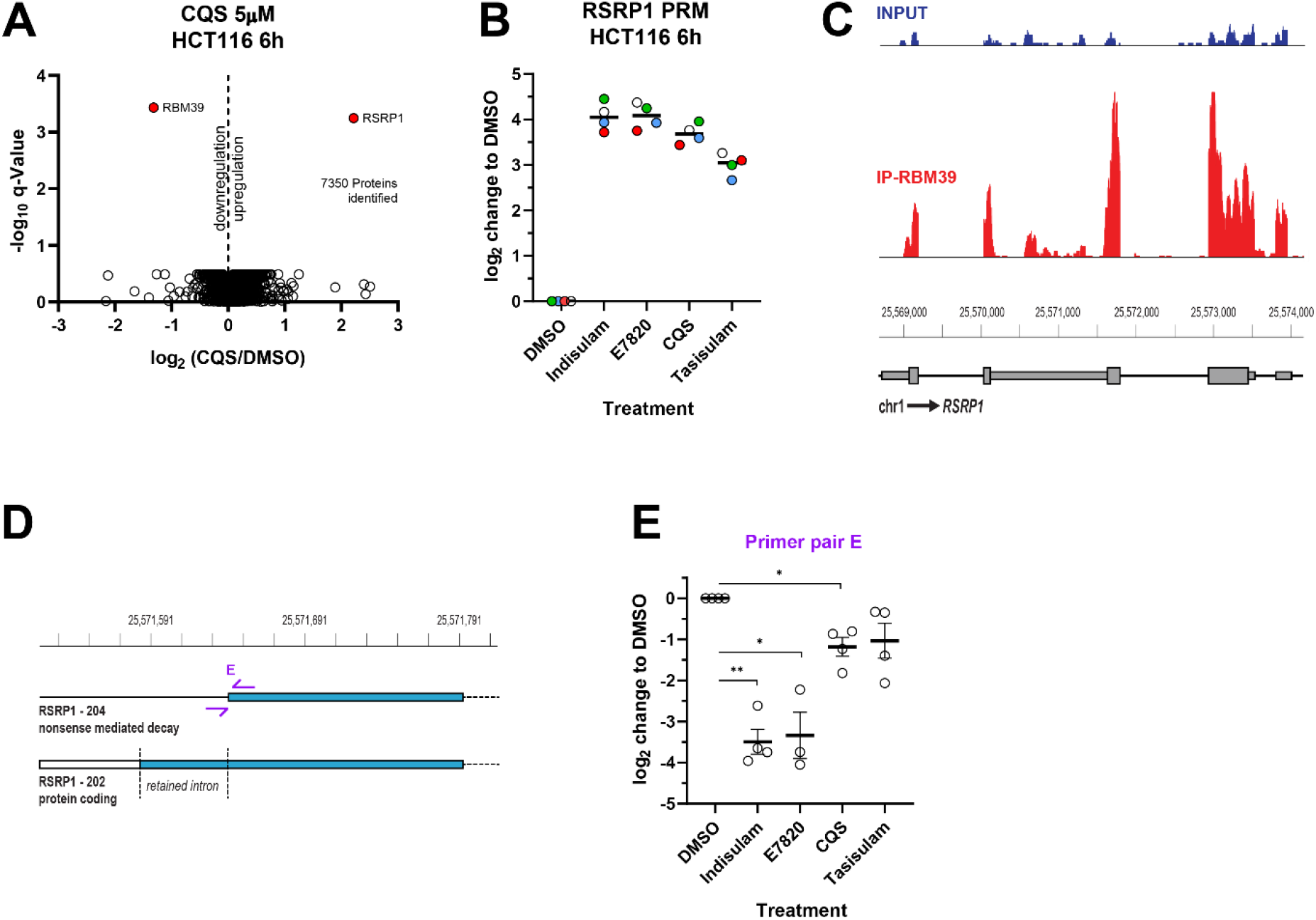
RSRP1 is upregulated by RBM39 degradation. **A)** Change in protein levels relative to vehicle treatment (DMSO) in HCT-116 cells treated with CQS at 5 μM for 6 h quantified as in Figure 2A. **B)** Change in RSRP-1 protein levels relative to vehicle treatment (DMSO) in HCT-116 cells treated with the sulfonamides at 5 μM for 24 h and quantified by taking the median of the level of 4 unique peptides (each individual peptide is shown as a different coloured dot) measured in a parallel reaction monitoring (PRM) experiment (see Figure S1D-E). **C)** eCLIP data shows strong enrichment relative to input of RSRP-1 transcripts after RBM39 immunoprecipitation. **D)** Comparison of non-coding RSRP1 transcript (202) and coding RSRP1 transcript (204) shows missing intron which was analyzed by primer pair E (full transcript list Figure S2A). **E)** Change in amplicon levels from primer pair E relative to vehicle treatment (DMSO) in HCT-116 cells treated with the sulfonamides at 5 μM for 6 h. Values represent mean −ΔΔ C_t_ values ±SEM from biological triplicates (*n = 3*, dots show individual values). Statistical analysis was performed using parametric unpaired t-test with Welch’s correction (*p < 0.0332, **p < 0.0021, ***p < 0.0002) (See Figure S2B).

### RSRP-1 splicing is altered by RBM39

RBM39 and RBM23 degradation caused by ArSulfs causes extensive splicing changes.[14, 21, 23] RBM23 degradation, however, has been shown to contribute little to ArSulf pharmacology.[21] Hence to identify the precise mechanism of ArSulf toxicity, we needed to identify those proteins most affected by RBM39 degradation. In our proteomics data we found the Arg/Ser-rich RSRP1 protein as a strongly upregulated protein (**Figure 2B & 3A**). Although RSRP1 likely has nothing to do with indisulam toxicity,[24] we decided to examine its regulation to build a workflow for characterizing RBM39 splicing targets. We hypothesized that RSRP1 might be a splicing target of RBM39, and that loss of RBM39 favors a particular coding transcript. In our hands commercial RSRP-1 antibodies performed poorly in Western blotting, so we validated RSRP-1 upregulation by parallel reaction monitoring (PRM) using four unique heavy peptides for RSRP-1 (**Figure 3B and Figure S1D-E**). Indeed, RSRP-1 was upregulated for each ArSulf in comparison to DMSO control (**Figure 3B**) in a DCAF15 dependent manner (**Figure S1D**). To validate that RSRP1 is a direct RBM39 target, we first analyzed the RBM39 eCLIP data and found that RSRP1 transcripts were significantly enriched by RBM39 immunoprecipitation (**Figure 3C**). To understand how RSRP1 was being regulated, we designed several primer pairs that could distinguish alternately spliced RSRP1 transcripts by qPCR (**Figure S2A** for the full set of primers used). Notably, we identified that a retained intron distinguished a coding transcript (RSRP1-202) from a transcript (RSRP1-204) leading to nonsense-mediated decay (NMD) (**Figure 3D**). We postulated that aberrant splicing due to low RBM39 levels leads to the retained intron and increases the abundance of coding-relative to the non-coding transcripts. According to the classification in the Ensembl database the region analyzed by Primer pair E was only part of non-coding RSRP-1 transcripts, such as RSRP1 −204 (**Figure 3D, Figure S2A**). Indeed, qPCR indicated that all the ArSulfs targeting RBM39 lead to down-regulation of this region (**Figure 3E**). These changes were also observed in K562 cells treated with ArSulfs (**Figure S1G**), but were absent in HCT116^DCAF15−/−^ control cells (**Figure S3H**). Taken together, these results support that RBM39 degradation changes alternative splicing decisions, ultimately causing up-regulation of the RSRP-1 protein. Although little is known about RSRP-1,[24] its low gene effect score (which measures the sensitivity of a cell line to CRISPR knockouts in a particular gene) [25] from the CRISPR avana dataset (see data at depmap.org)[26] suggests its upregulation may not be a significant contributor to ArSulf toxicity. Nevertheless, our experimental approach with RSRP1 suggested that integration of eCLIP datasets could help to identify transcriptional effects in proteomics data with ArSulfs.

### Splicing of kinesin proteins KIF20A and KIF20B is altered upon RBM39 degradation

Inspired by the RSRP-1 findings, we wondered whether there might be a more general way to identify direct transcriptional targets of RBM39 degradation. Hence, we overlayed the eCLIP targets for RBM39[23] with proteins identified in TMT proteomics datasets after ArSulf treatment. In this way we could find proteins whose levels changed significantly upon ArSulf treatment, and whose transcripts seemed to have a direct interaction with RBM39. The kinesin motor proteins KIF20A and KIF20B were just such cases: consistently downregulated after exposure to the ArSulfs and highly enriched in the eCLIP data (**Figure 4A, Figure S1F**). We decided to take a closer look at both proteins. Upon analyzing the four transcripts of KIF20B listed in the Ensembl database, we found that three of them were coding and the other one retained an intron, making it non-coding. We hypothesized that RBM39 loss was causing retention of this intron, leading to a preference for the non-coding transcript (**Figure 4B**). Indeed, qPCR analysis showed a significantly higher retention of the intron in ArSulf treated HCT116 and K562 cells (**Figure 4D, Figure S2D**), along with a slight downregulation of the coding transcripts (**Figure 4C, Figure S2A**).

**Figure 4.**
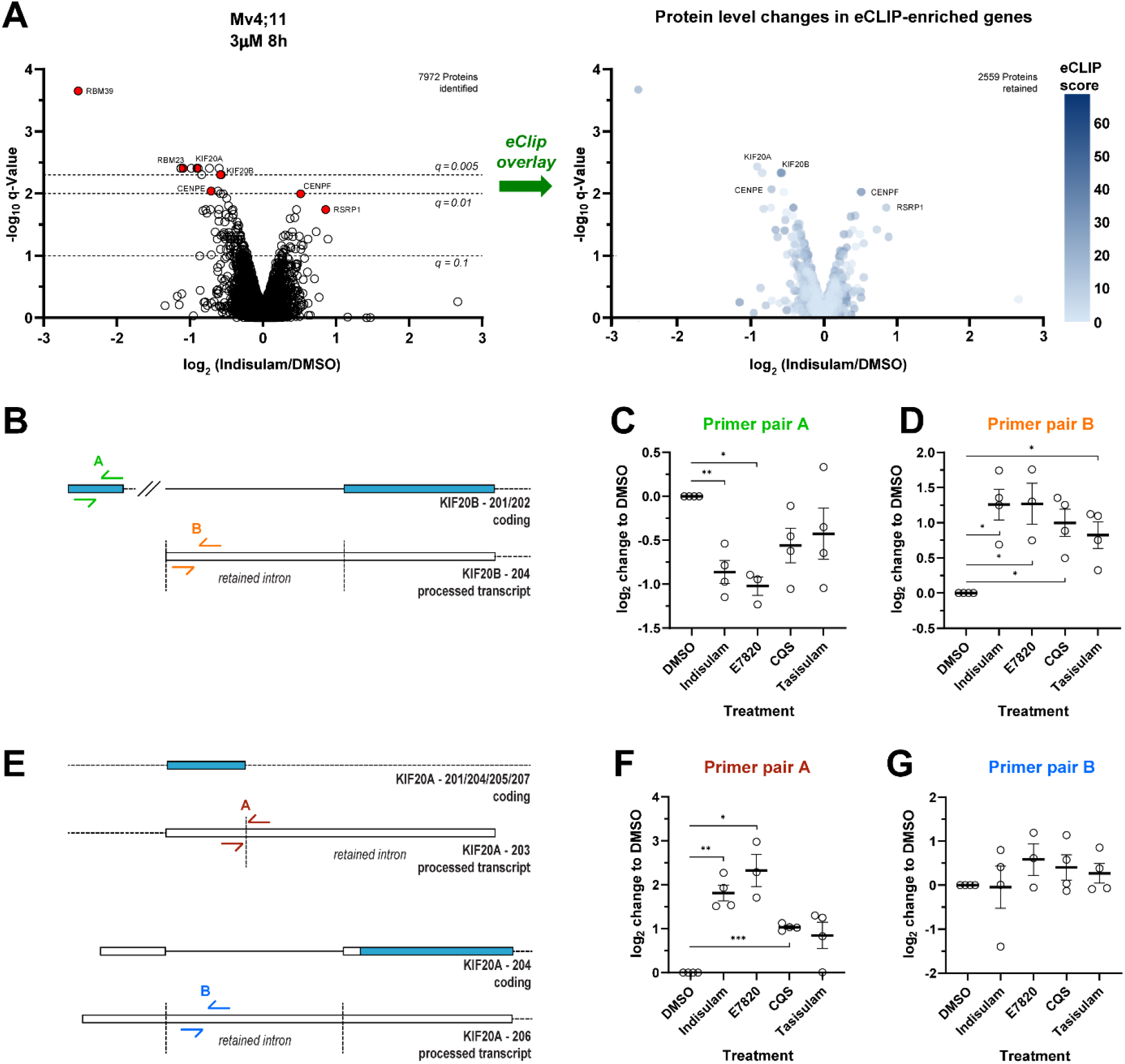
KIF20A and KIF20B are downregulated by RBM39 degradation. **A)** Plot of proteins changes quantified from proteomics as in Figure 2A) with their scores from RBM39 eCLIP enrichment (as colors). As shown in Figure 2A) proteomics was done in Mv4;11 cells treated with Indisulam at 3 μM for 8 h. Strongly up- or down-regulated sequences with high eCLIP scores are likely to be transcriptional targets of RBM39, such as KIF20A or KIF20B (see Figure S1F). **B)** Comparison of noncoding KIF20B transcript (204) and coding KIF20B transcript (201) shows retained intron which was analyzed by primer pair B, whereby the coding transcript was analyzed by primer pair A (full transcript list Figure S2A). RBM39 loss leads to intron retention and preferential formation of a non-coding transcript in KIF20B splicing. **C)** Change in amplicon levels from primer pair a relative to vehicle treatment (DMSO) in HCT-116 cells treated with the sulfonamides at 5 μM for 6 h. Values represent mean −ΔΔ C_t_ values ±SEM from biological triplicates (n = 3, dots show individual values). Statistical analysis was performed using parametric unpaired t-test with Welch’s correction (*p < 0.0332, **p < 0.0021, ***p < 0.0002) (see Figure S2B). **D)** As in C) for primer pair B. **E)** As in B) comparison of the coding transcript and non-coding transcripts show retained introns which were analyzed by primer pair A and B (full transcript list Fig S2A). RBM39 loss leads to intron retention and preferential formation of a non-coding KIF20A transcript after splicing. **F)** As in C) for primer pair A. **G)** As in C) for primer pair B.

KIF20A has 7 transcripts, of which only 4 are coding. Two of the remaining non-coding transcripts (KIF20A-203, KIF20A-206) retain an intron and the last one skips an exon, leading to NMD. We decided to analyze these transcripts (KIF20A-203 and KIF20A-206) more closely in the region of the retained introns (**Figure 4E**). qPCR with corresponding primers identified KIF20A-203 as being upregulated, while the other was unaffected in both HCT116 and K562 cells (**Figure 4F-G, Figure S2A, Figure S2D**). Importantly, the changes in both KIF20B and KIF20A are phenocopied by siRNA mediated RBM39 knockdown in HCT116 cells (**Figure S2C**) and are absent upon ArSulf treatments of HCT116^DCAF15−/−^ (**Figure S2E**) cells as well as in HCT116 cells where RBM23 is knocked-down with an siRNA (**Figure S2C**). Collectively, these results show that RBM39 directly mediates KIF20A and KIF20B splicing and that RBM39’s degradation causes aberrant splicing that ultimately alters the protein levels of both kinesins. KIF20A and KIF20B changes seemed likely contributors to ArSulf toxicity since these are important mitotic proteins often up-regulated in cancer.[27–30] In summary, by combining proteomics and eCLIP analysis, we identified RSRP-1, KIF20A and KIF20B as transcriptional targets of RBM39 that are strongly affected at the protein level, providing a workflow that broadly explores the proteome impact of targeting splicing factors.

### ArSulf treatment phenocopies KIF20A and KIF20B down-regulation

Various kinesin motor proteins play an essential role across each stage of cell division[31–34] and are up-regulated in nearly all types of cancer. [35] Whole proteome screens have revealed that KIF20A and KIF20B are upregulated in mitosis[36–40], most likely in the G2/M phase and cytokinesis/telophase transitions.[41] Both KIF20A (a.k.a. MKLP2) and KIF20B (a.k.a. MPP1) belong to the Kinesin-6 family and are involved in the organization and regulation of the mitotic spindle and cytokinesis.[33, 42] Inhibiting KIF20A with the small molecule paprotrain or with an siRNA has resulted in varying effects on cells,[43] leading to attenuated cell growth or apoptosis.[44] A consistent finding, however, is that inhibiting KIF20A and KIF20B activity leads to failure in cytokinesis and multinucleated cells.[33, 45–48] If ArSulf treatment was causing toxicity by altering kinesin splicing, we should also observe increases in multi-nucleated cells upon treatment. Indeed, exposure for 24h resulted in a measurable increase in multinucleated cells, but the effect after 48h was more dramatic (**Figure 5A**), with 20% of cells showing multinucleation. As would be expected in cells experiencing such mitotic stress, the 48h time-points also showed high cytotoxicity (**Figure 5B, Figure S3A**). Additionally, cells remaining after 48h showed significantly reduced mitosis (**Figure 5B**). In summary, lowering of KIF20A and KIF20B protein levels after pharmacological RBM39 degradation is a feature we have observed in five cell lines covering three indisulam-sensitive cancer lines originating from different organ systems (hepatocellular carcinoma (HCT116), leukemia (Mv4;11, MOLM-13, KBM7 (data from others)[49]), and neuroblastoma (IMR-32 (data from others)[12]). We further demonstrate that multinucleation and cytotoxicity are downstream effects of these changes in HCT116 – a cell line particularly sensitive to ArSulfs.

**Figure 5.**
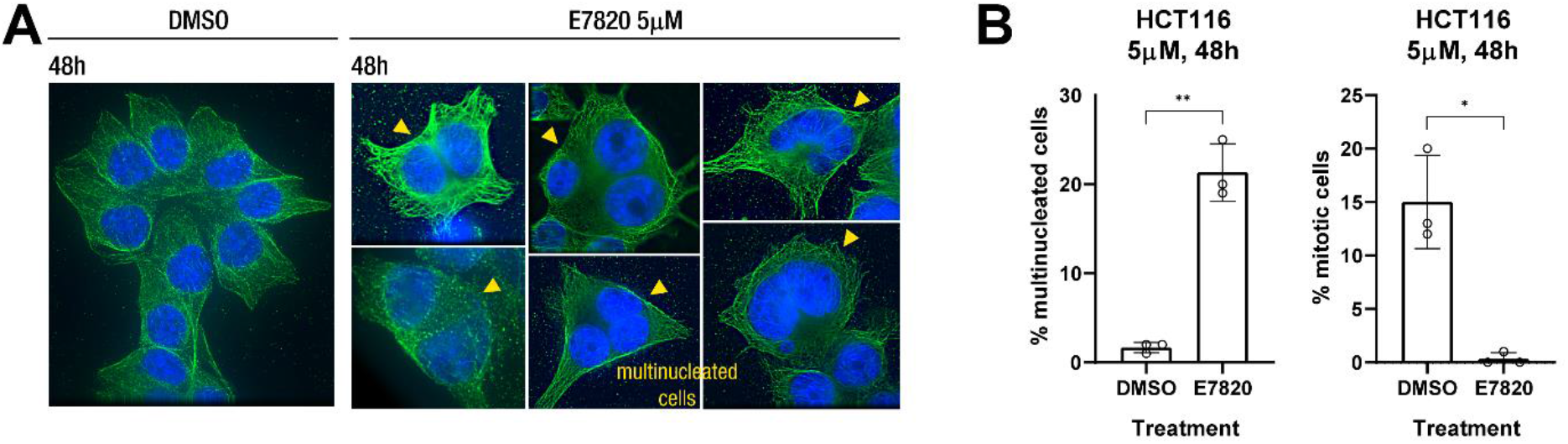
KIF20A and KIF20B downregulation leads to multinucleated cells. **A)** Immunofluorescence images of HCT-116 cells treated with DMSO or E7820 (5 μM) for 48h and stained for DNA (blue), α-Tubulin (green). **B)** Amount of multinucleated and mitotic cells observed in vehicle treated (DMSO) and E7820 treated HCT-116 cells at 5 μM for 48 h. Values represent mean values ±SD from counting 100 cells per treatment in biological triplicates (*n = 3*, dots show individual values). Statistical analysis was performed using parametric unpaired t-test with Welch’s correction (*p < 0.0332, **p < 0.0021, ***p < 0.0002). Example images showed In Figure S3A.

### Effect of RBM39 degradation on cell cycle progression

Although downregulation of the targets of the HOXA9 transcription factor (such as MCL1) has been identified by functional genomics as a potentially important contributor to ArSulf activity in leukemia,[23] the major contributors to activity in other cancer types are unknown – although a recent preprint has identified potential contributors in neuroblastoma.[12] To assess the cellular impact from the individual effects caused by ArSulfs (i.e. RBM39 and RBM23 downregulation and DCAF15 inhibition) we did TMT-proteomics in cells with siRNA knockdown of each gene (**Figure S3B**). Neither RBM23, nor DCAF15 knockdown caused big changes in the proteome. However, applying a STRING network analysis on the most significantly altered proteins in cells with RBM39 knockdown and in ArSulf treated cells predominantly revealed processes involved in the cell cycle or mitosis (**Figure S3C-D**). Doing a pathway analysis (using Enrichr) on the most enriched genes in the eCLIP dataset (**Supplementary Data Table 1**) reassured this observation. Additionally, when searching for molecules whose toxicity profile correlated most strongly with indisulam using the PRISM dataset (data downloaded from depmap.org)[50–52] we found that indisulam-mediated cell death strongly correlates with other mitotic poisons such as tubulin disruptors, as well as inhibitors of aurora kinase, polo-like kinase inhibitors, and mitotic kinesins (**Figure 6A**). In contrast, no correlation was found upon analysis for molecules that are known to cause G1 cycle arrest, such as mTOR, EGFR, or CDK4 inhibitors (see **Supplementary Data Table 2**). Hence, ArSulf treatment seems to cause a mitotic poison phenotype through a mechanism distinct from most mitotic poisons.

**Figure 6.**
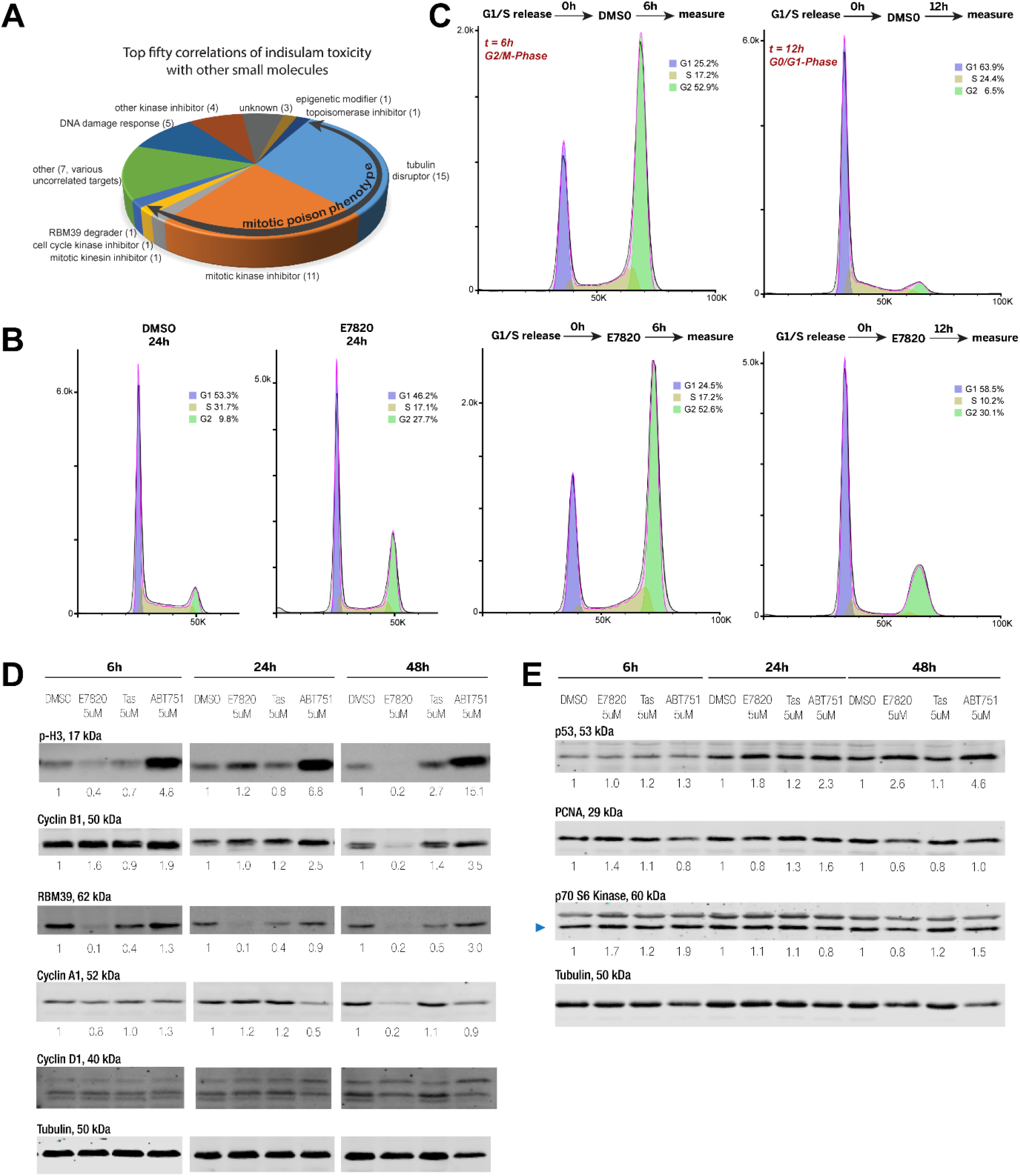
RBM39 degradation causes changes in the cell cycle. **A)** The PRISM dataset reveals that indisulam toxicity across hundreds of cancer cell lines correlates with the toxicity profile observed with mitotic poisons. **B-C)** Cell cycle was determined using flow cytometry. The data was analyzed using FlowJo 10.5.3 and the population quantified using the Watson (pragmatic) model. **B)** Asynchronous cells were treated with DMSO or E7820 (5 μM) for 24h or 48h. **C)** Cells were synchronized in G1/S using a double thymidine block and treated with DMSO or E7820 (5 μM) upon release and treated for 6h or 12h (see Figure S4B). **D)** Western Blot of Anti-Phospho-Histone H3 (Ser10), Anti-Cyclin A1, Anti-Cyclin B1, Anti-Cylcin D1, Anti-RBM39 and Anti-Tubulin of HCT-116 cells treated with vehicle (DMSO), E7820 (5 μM), Tasisulam (5 μM), ABT-751 (5 μM) for 6h (left), 24h (middle) or 48h (right). The numbers on the Blot indicate the signal intensities of the bands relative to the Tubulin bands, normalized to DMSO treatment as quantified with Image Studio Lite. **E)** As in D) but Western Blot of Anti-p53, Anti-PCNA, Anti-p70S6 and Anti-Tubulin.

Next, we took a closer look at the cell cycle effects. In particular, we used flow cytometry to explore the effects of ArSulf exposure on cell cycle progression in HCT116 cells. We choose E7820 for these experiments since our proteomics and PRM data suggest it is the most potent ArSulf. In asynchronous cells, E7820 treatment after 24h leads to a near tripling of cells (9.8%➔27.7%) in G2/M (**Figure 6 B**). In HCT-116 cells synchronized in the S-phase we observed that after 6h the phases of the E7820 and DMSO treated cells were indistinguishable (**Figure 6C**), although Western blotting confirmed RBM39 degradation (**Figure S4C**). After 12h, however, the DMSO cells had completed a second cell cycle while the E7820 cells still had a substantial fraction at G2/M (**Figure 6C, Figure S4B**). Consistent with the immunofluorescence data (see **Figure 5** and associated discussion), at 48h we observed high toxicity in the E7820 sample (**Figure S4A**). The latency period for toxicity is consistent with the mechanism of action of indisulam, since rapid RBM39 loss is then followed by splicing changes and slow proteome adaption. These results suggest that E7820 does not impact the cell cycle immediately upon RBM39 degradation, but on second or subsequent cycles G2/M arrest is observed, ultimately leading to multinucleation or cell death.

To explore further the impact on mitosis we compared the effect of E7820 to the known microtubule inhibitors nocodazole, ABT-751 as well as the CENPE inhibitor GSK-923295. These molecules all cause a stronger G2/M arrest than ArSulfs (**Figure 6 D-E, Figure S5C**), but proteomics data reveals that, in contrast to ArSulfs, they all cause an increase in KIF20A and KIF20B levels (**Figure S5A**). Mitotic markers cyclin B and phosphorylated histone H3 (Ser10), as well as cyclin A, showed moderate increases at early time points (6h and 24h), but at 48h all of these markers drop (**Figure 6D, Figure S5C**) (control experiments in HCT-116^DCAF15−/−^ cells confirm the involvement of DCAF15 (**Figure S5D**)). Furthermore, proliferation markers PCNA and p70S6K were also downregulated in the surviving cells (**Figure 6E**), consistent with these cells having suffered failures in cell division. Taken together these results suggest that the ArSulf E7820 first leads to a G2/M arrest, followed by cell death or multi-nucleation due to mitotic failure. The results with E7820 contrast sharply with the other mitotic inhibitors; for example, ABT-751 gives a strong and durable G2/M arrest, with little change in markers over 48h (**Figure 6D-E**).

## Discussion

### Molecular glues can drug difficult-to-target proteins

The *de novo* discovery of new molecular glues that target a specific class of molecules is a challenging screening endeavor. Until recently, all monovalent molecular glues had come from either natural selection, phenotypic screening, or happenstance.[53–55] The IMiDs, exemplified by lenalidomide and pomalidomide, are continuing to deliver new hits against transcription factors, a protein class often dubbed ‘undruggable’ by conventional means. Using the IMiD scaffold’s binding to the E3 ligase substrate receptor protein CRBN as a molecular anchor, IMiDs have also enabled the creation of bifunctional molecules where E3 ligase substrate selection can be modularly reprogrammed (PROTACs). The saga of the IMiDs is what makes the discovery of the ArSulfs so tantalizing: these bind an E3 ligase substrate receptor, DCAF15, and alter its substrate selection to degrade the splicing factor RBM39. Could ArSulf targeting of splicing factors replicate the story of the IMiDs targeting zinc-finger transcription factors? RNA binding proteins are also a large and largely ‘undruggable’ class of proteins that bind and regulate macromolecular assemblies. Modulating alternative splicing of disease-causing proteins[56] or targeting splicing mutated cancers[57] is an emerging area of therapeutic development and the ArSulfs represent an excellent starting point because they profoundly alter splicing decisions. Although directing ArSulfs to target other RNA-binding proteins is an unmet challenge, our results outline a strategy for quickly assessing consequences of RBM39 degradation. With this analysis workflow in hand, it would be interesting to examine ArSulfs in diseases where targeting faulty splicing[58] or inducing exon skipping[59, 60] have delivered therapeutics.

### ArSulfs degrade only the RBM39/RBM23 RRM2 motif

We find the second RRM motif (RRM2) found in RBM39 to be a strict degron in terms of primary protein sequence. This finding holds true across several cell lines and ArSulf derivatives. Our current model for why DCAF15 is not amenable to reprogramming is that once RBM39 is depleted the poor affinity of ArSulfs to DCAF15 is not sufficient to support active ternary complex assembly with low affinity RRMs, a conclusion supported by data from others.[17] A tight-binding small molecule for DCAF15 is therefore an important future goal for the development of DCAF15-dependent PROTACs and molecular glues.

### Distinguishing transcriptional and post-transcriptional effects of ArSulfs

Broadly distinguishing transcriptional from post-transcriptional effects will be key for deploying DCAF15 against new proteins and for understanding how RBM39 degradation exerts its effect on cells. These requirements are especially important for ArSulfs, since they degrade a splicing factor, leading to widespread transcriptional changes. Comparing eCLIP and proteomics data helped us disentangle transcriptional from molecular glue effects proteome-wide, a workflow that helped us confirm RBM23 as a molecular glue target of the ArSulfs and to identify RSRP-1, KIF20A, and KIF20B as splicing substrates of RBM39 that lead to strong changes at the protein-level. We believe this data analysis workflow will be immediately helpful for finding molecular glues targeted to DNA or RNA-binding proteins that have downstream effects on transcription, such as transcription factors or splicing factors. Projecting forward from our findings, SLAM-seq[61] coupled with proteomics would offer a global comparison of transcriptome/proteome changes for any target protein in response to treatment with protein degrading molecules.

### RBM39 degradation causes strong changes in cell cycle and mitosis, likely due to KIF20A and KIF20B downregulation

Although there are certain to be lineage specific impacts of changes in splicing programs,[23, 62] there may also be specific ArSulf-induced splicing changes that are broadly toxic. We find in several different ArSulf-sensitive cancer cell lines that KIF20A and KIF20B is strongly down-regulated at the protein level. Cell cycle analysis, on the other hand, shows that a larger proportion of cells treated with ArSulfs remain in G2/M. Given that KIF20A and KIF20B are typically up-regulated in G2/M (see **Figure S5A** for proteomics data with molecules that cause G2/M arrest through three different mechanisms: nocodozole, ABT-751, and GSK-923295 – all of which show KIF20A and KIF20B increases) these observations are best explained by regulation of these mitotic kinesins by RBM39. eCLIP data as well as transcript analysis show that changes in splicing programs are consistent with a direct physical interaction between RBM39 and KIF20A and KIF20B transcripts. Such dramatic changes in mitotic kinesins, which are essential for proper mitosis, would suggest a mitotic poison phenotype of the ArSulfs. Indeed, cell cycle and cellular features after ArSulf treatment are consistent with a mitotic poison, for example increases in G2/M (**Figure 6B**) and multinucleation in surviving cells (**Figure 5A-B**). Examination of how indisulam toxicity correlates with that of other cancer therapeutics uncovers a striking correlation to other mitotic poisons (**Figure 6A**), with tubulin disruptors, mitotic kinases, and kinesin inhibitors dominating the top correlations. Although tubulin disruptors are a mainstay of cancer treatment, they invariably come with strong nervous system toxicity.[63] Targeting other mitotic proteins (for example inhibitors of polo-like kinases, aurora kinases, and mitotic kinesins) might recapitulate the strong cell-killing effect of tubulin disruptors while reducing toxicity, but so far these have largely failed. As such, novel mechanisms that disrupt mitosis could provide exciting starting points for drug development. RBM39’s ability to target multiple mitotic kinesins would seem to be a unique avenue for disrupting mitosis.

## Supporting Figures

**Figure S1.**
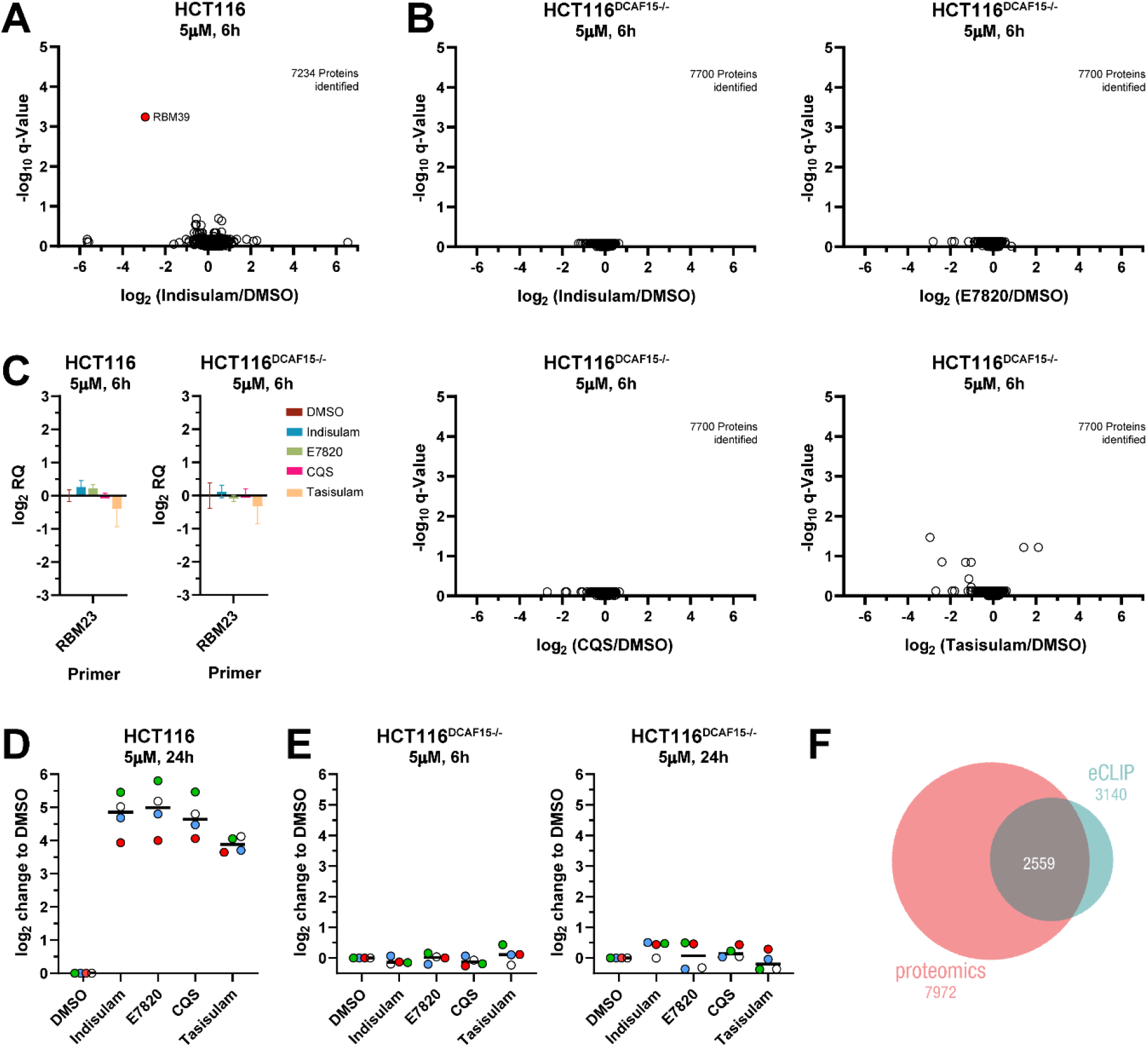
**A)** Change in protein levels relative to vehicle treatment (DMSO) in HCT-116 cells treated with the sulfonamides at 5 μM for 6 h quantified by TMT labelling and LC MS/MS analysis, versus q-Value (Benjamini–Hochberg corrected p-Value from Bayes moderated t-statistics). **B)** As for A) but in HCT-116^DCAF15−/−^ cells. **C)** qRT-PCR triplicate measurement showing the change in amplicon levels of RBM23 relative to vehicle treatment (DMSO) in HCT-116 cells treated with the sulfonamides at 5 μM for 6 h The quantification was done by using StepOne Software v2.3 to calculate RQ (same as 2^−*ΔΔCt*^) relative to DMSO using GAPDH as the housekeeping gene. The error bars were calculated from RQMax and RQMin as defined in the software using 95% CI. **D)** Change in RSRP-1 protein levels relative to vehicle treatment (DMSO) in HCT-116 cells treated with the sulfonamides at 5 μM for 24 h and quantified by taking the median of the level of 4 unique peptides measured in a PRM experiment **E)** As for C) but in HCT-116^DCAF15−/−^ cells treated for 6 h (left) and 24 h (right). **F)** Venn diagram showing overlap between the genes identified in proteomics datasets and eCLIP dataset used in Figure 4A).

**Figure S2.**
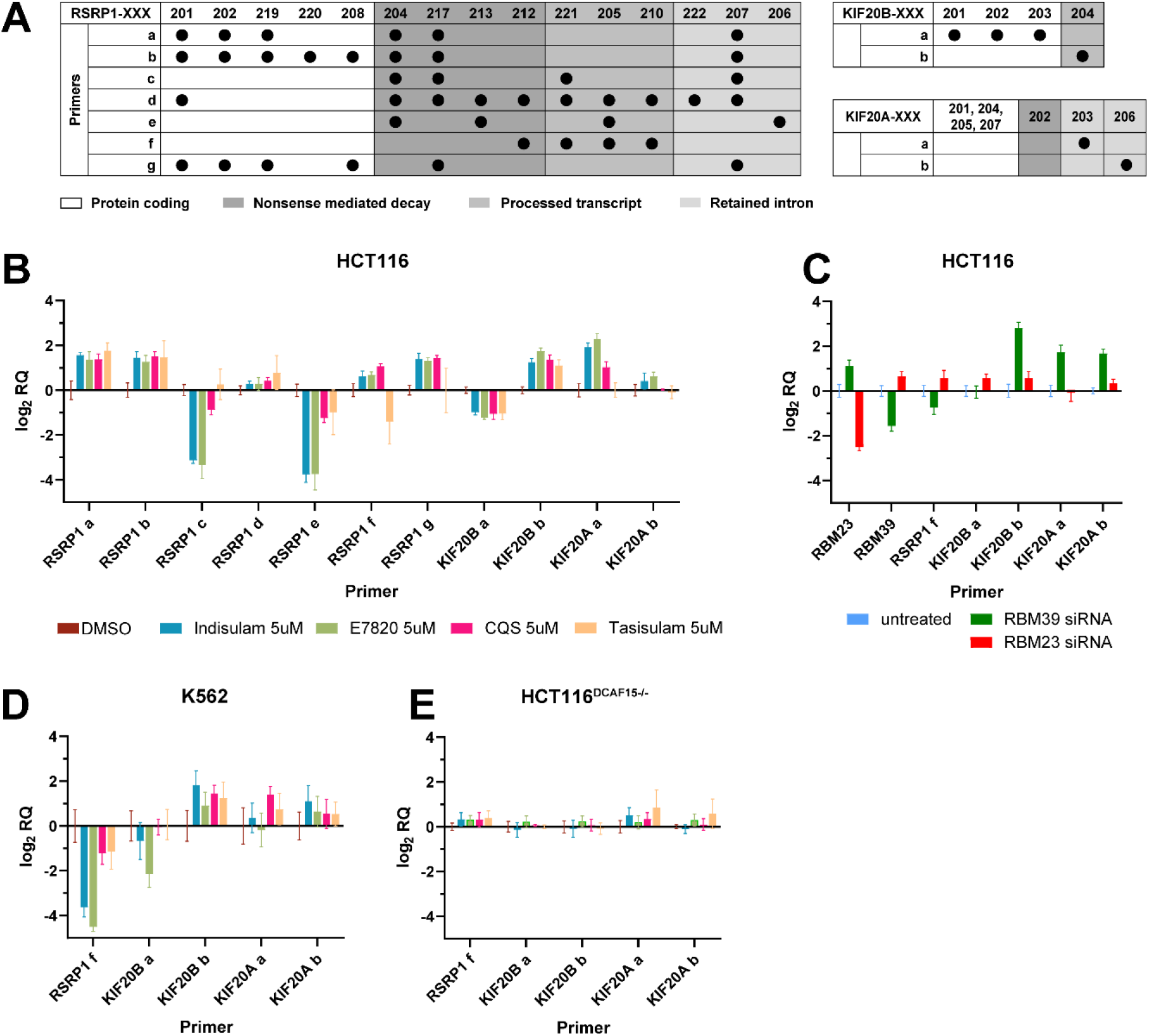
**A)** Summary of all primers used for qRT-PCR and the transcripts of RSRP1 and KIF20B that contain the amplified amplicon. **B)** Example of a qRT-PCR triplicate measurement showing the change in amplicon levels of the genes relative to vehicle treatment (DMSO) in HCT-116 cells treated with the sulfonamides at 5 μM for 6 h. The quantification was done as in Figure S1C). **C)** Amplicon levels in siRNA mediated RBM39 knocked down HCT-116 cells quantified as in B). **D)** As for B) but in K562 cells **E)** As for B) but in HCT-116^DCAF15−/−^ cells.

**Figure S3.**
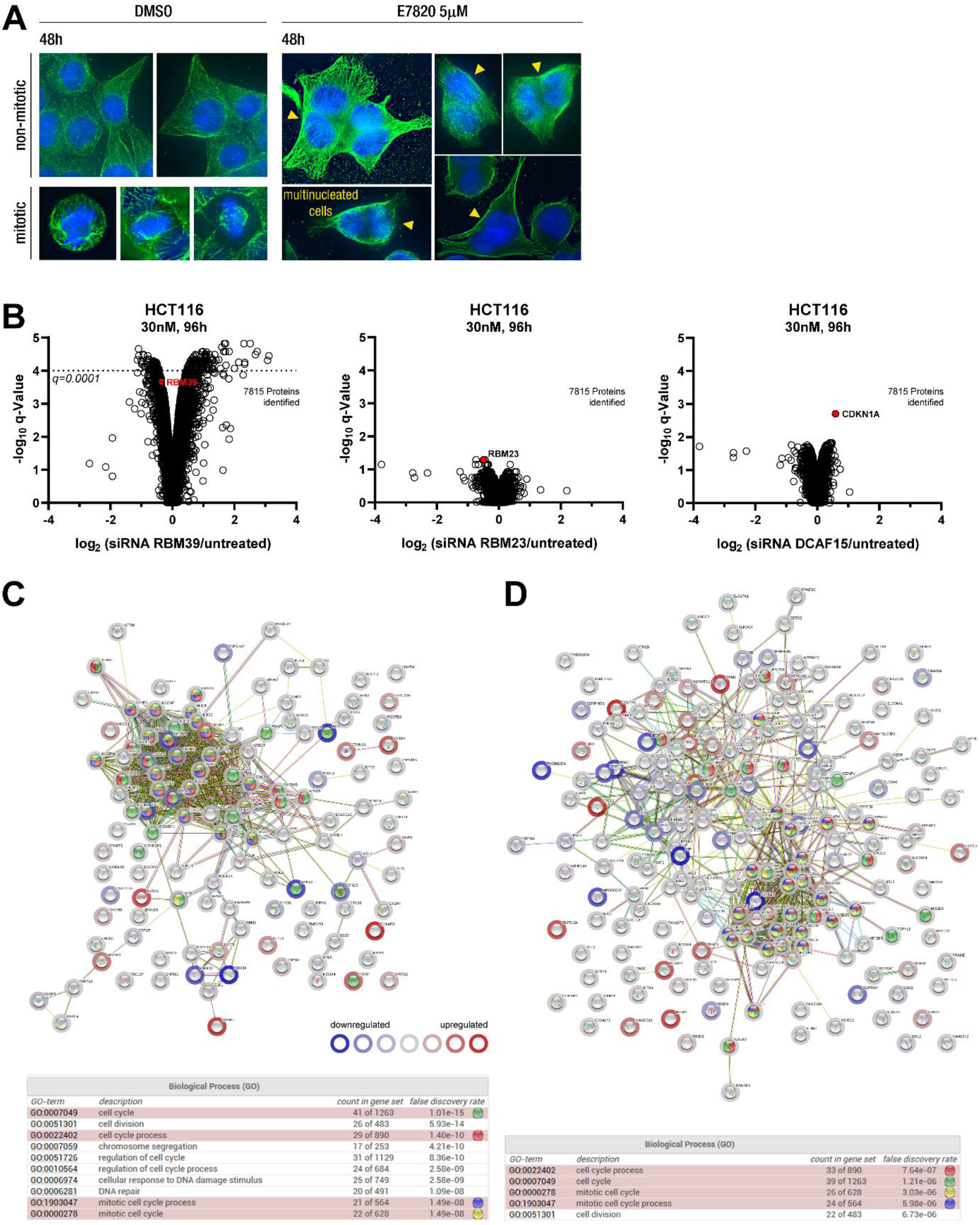
**A)** Example of immunofluorescence images of HCT-116 cells treated with DMSO or E7820 (5 μM) for 48h and stained for DNA (blue), α-Tubulin (green) used to quantify mitotic and multi-nucleated cells for Figure 5B). B-C) Relative protein levels between untreated HCT-116 cells and cells treated with siRNA (30 nM) against RBM39, RBM23 and DCAF15 for 96h. Proteins levels were measured and analysed as for Figure S1A). **C-D)** STRING network analysis of significantly regulated proteins, ranked according to the extent of regulation, in cells upon RBM39 downregulation. Network analysis of proteins regulated in **C)** Mv4;11 cells treated with Indisulam (3 μM) for 8h (Figure 2A) with q<0.1, **D)** HCT-116 cells treated with RBM39 siRNA (30 nM) for 96h with q<0.0001 (Figure S3B). Ball color indicates biological process and ring color indicates extent of up (red) or down (blue) regulation.

**Figure S4.**
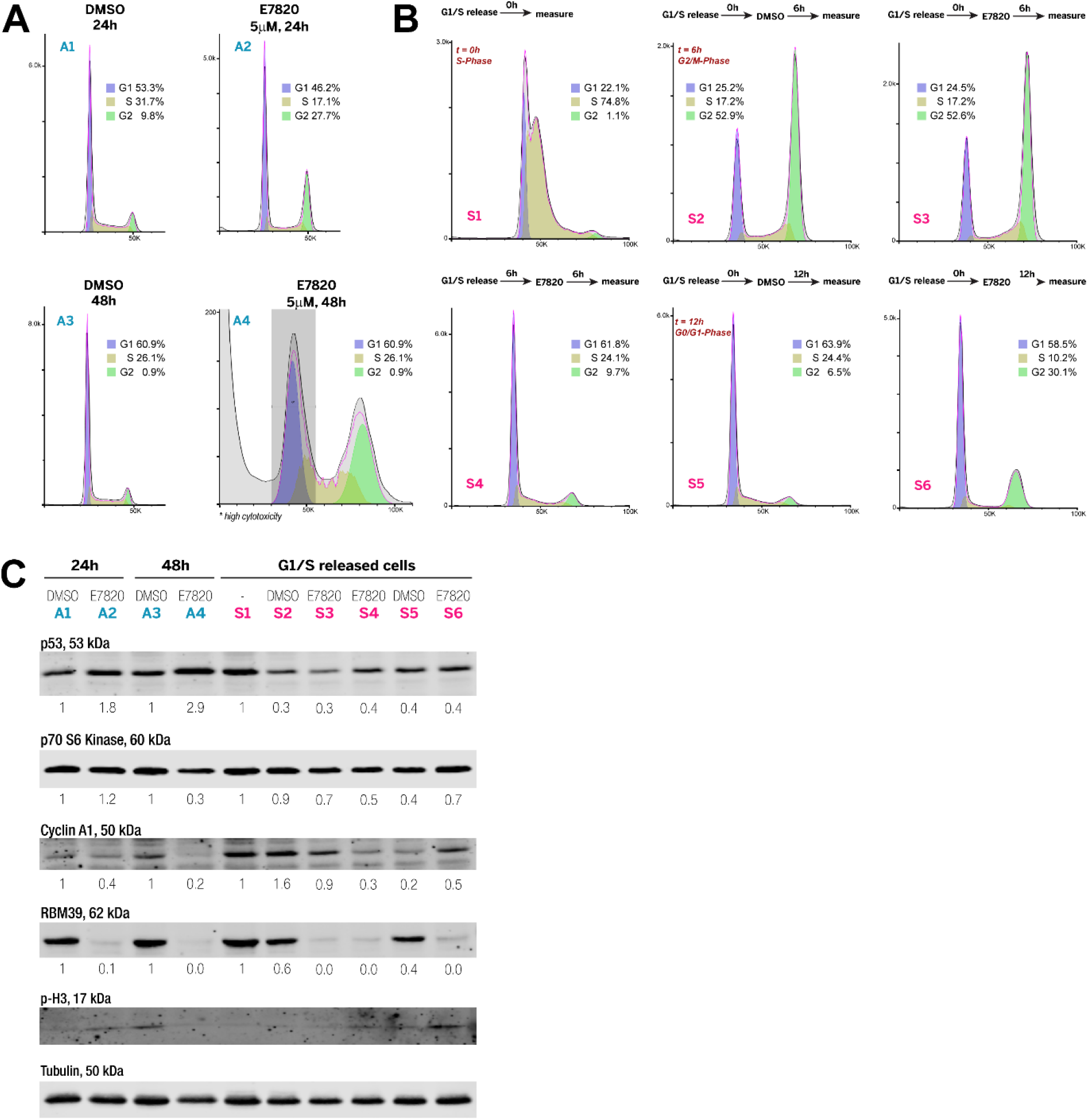
E7820 impacts cell cycle in HCT-116 cells. **A-B)** Cell cycle was determined using flow cytometry. The data was analyzed using FlowJo 10.5.3 and the population quantified using the Watson (pragmatic) model. **A)** Asynchronous cells were treated with DMSO or E7820 (5 μM) for 24h or 48h. **B)** Cells were synchronized in G1/S using a double thymidine block and treated with DMSO or E7820 (5 μM) at different time points after release (0h or 6h) and treated for 6h or 12h. t indicates the time after release. **C)** Western Blot of Anti-Phospho-Histone H3 (Ser10), Anti-Cyclin A1, Anti-RBM39, Anti-p53, Anti-PCNA, Anti-p70S6 and Anti-Tubulin of the HCT-116 cells analyzed in A) and B). The numbers on the Blot indicate the signal intensities relative to the Tubulin bands, normalized to DMSO treatment as quantified with Image Studio Lite.

**Figure S5.**
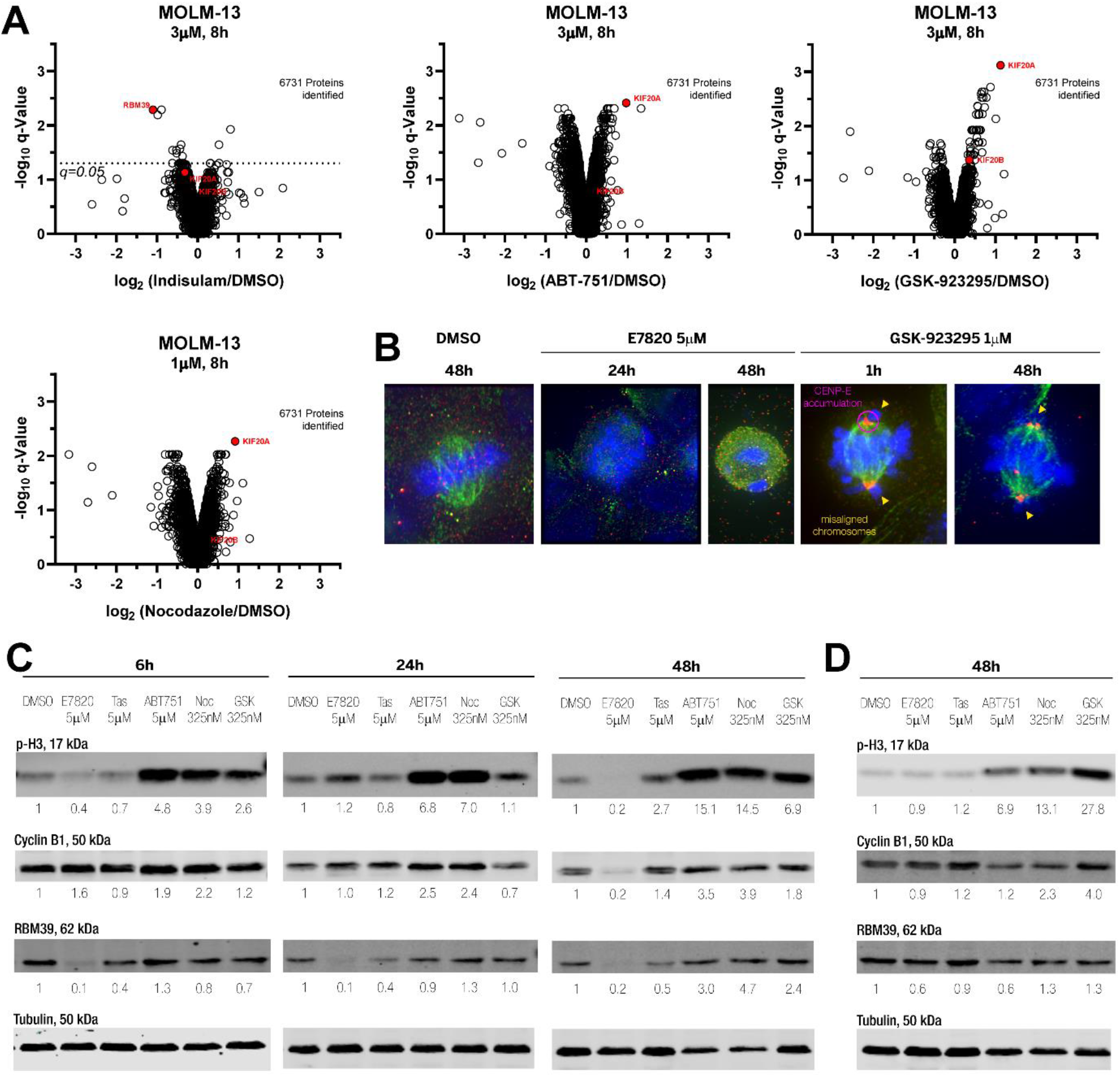
**A)** Relative protein levels between untreated MOLM-13 cells and cells treated with Indisulam (3 μM), ABT-751(3 μM), GSK-923296 (3 μM) and Nocodazole (1 μM) for 8h. Proteins levels were measured and analysed as for Figure S1A). **B)** Immunofluorescence images of HCT-116 cells treated with DMSO, E7820 (5 μM) or the CENP-E (1 μM) inhibitor GSK-923295 for the indicated times and stained for DNA (blue), CENPE (red) and a-Tubulin (green). **C)** Western Blot of Anti-Phospho-Histone H3 (Ser10), Anti-Cyclin B1, Anti-RBM39 and Anti-Tubulin of HCT-116 cells treated with vehicle (DMSO), E7820 (5 μM), Tasisulam (5 μM), ABT-751 (5 μM), Nocodazole (325 nM) or GSK-923295 (325 nM) for 6h (right), 24h (middle) or 48h (left). The numbers on the Blot indicate the signal intensities of the RBM39 bands relative to the Tubulin bands, normalized to DMSO treatment as quantified with Image Studio Lite. **D)** As in A) but in HCT-116^DCAF15−/−^ cells for 48h.

## Experimental

### Cell lines and cell culture

The adherent HCT-116 (a kind gift from M. Hall, University Basel) cells were grown in DMEM with 10% FCS and 1% penicillin/ streptomycin. The human leukemia cells Mv4;11, MOLM-13, K-562 (kind gifts from J. Schwaller, University Basel) were grown in RPMI with 10% FCS and 1% penicillin/ streptomycin. The cells were kept in the an incubator at 37 °C with 5% CO_2_. The HCT-116^DCAF15−/−^ cells were generated as previously described.[14]

### Cell treatments

All cell treatments were done in standard 12 well plates. HCT-116 cells were seeded at 170k cells per well and K562, Mv4;11, MOLM-13 cells at a density of 250k cells per well. After 1 d, the cells were treated with compounds for the indicated period of time. The adherent cells were washed thrice with ice cold PBS after which they were scraped from the wells and centrifuged at 8.5k *g* for 15 min at 4 °C. The suspension cells were collected by centrifugation (12k g for 5 min), and washed by resuspending them in cold PBS and collectinon by centrifugation (12k g for 5 min) three times. The PBS was removed and the cell pellets were stored at −80 °C and either used for LC-MS analysis or Western blotting.

The compounds ABT-751 (*MedKoo Biosciences, Inc*.), Nocodazole (*MedChem Express*) and GSK-923295 (*MedChem Express*) were purchased.

The cells for RNA extraction and qPCR were not washed but directly collected in 1 mL TRI Reagent^®^ (TR 118) and stored at −80 °C.

### Western blot

The cell pellets were lysed in 50 μL lysis buffer (RIPA buffer, supplemented with Roche complete™ protease inhibitor and 100 mM orthovanadate). The lysate was incubated on ice over 30 min with periodic vortexing and subsequently centrifuged at 16.9k *g* for 15 min at 4 °C. Protein concentrations were measured using DC Protein Assay. The lysates were incubated in loading buffer (5x, SDS, DTT, bromophenol blue, glycerol) at 98 °C for 5 min. The lysates (15 μg per lane) were seperated on a 8% or 11% Tris-Glycine SDS-polyacrylamide gel and then transferred on nitrocellulose membranes (Amersham Protran 0.45 NC) using a discontinuos CAPS buffer system with the BioRad Trans-Blot Turbo System. The blots were blocked using Odyssey Blocking Buffer (TBS) and probed wiith antibodies for RBM39 (SigmaAldrich, HPA001591, 1:5k dilution), *α*-Tubulin (Abcam, ab7291, 1:10k dilution), p70 S6 kinase (SCBT, sc-8418, 1:500 dilution), PCNA (SCBT, sc-56, 1:500 dilution), p53 (SCBT, sc-126, 1:500 dilution), Cyclin A1 (SCBT, sc-271682, 1:500 dilution), Cyclin D1 (SCBT, sc-8396, 1:500 dilution), p-H3 (CST, 3377T, 1:1k dilution), Cyclin-B1 (GeneTex, GTX100911, 1:500). The blots were further developed using anti-Mouse (Licor, IRDye^®^ 680RD or 800CW Goat anti-Mouse IgG, 1:10k dilution) or anti-Rabbit secondary antiboy (Licor, IRDye^®^ 680RD or 800CW Goat anti-Rabbit IgG, 1:10k dilution) and the bands were visualized using Licor Odyssey CLx imager. The images were processed and the bands quantified using Licor Image Studio software.

### Sample preparation for TMT and PRM LC-MS/MS analysis

Cells were lysed in 50 μL lysis buffer (1% sodium deoxycholate (SDC), 0.1 M TRIS, 10 mM TCEP, pH = 8.5) using strong ultra-sonication (10 cycles, Bioruptor, Diagnode). Protein concentration was determined by BCA assay (Thermo Fisher Scientific) using a small sample aliquot. Sample aliquots containing 50 μg of total proteins were reduced for 10 min at 95 °C and alkylated at 15 mM chloroacetamide for 30 min at 37 °C. Proteins were digested by incubation with sequencing-grade modified trypsin (1/50, w/w; Promega, Madison, Wisconsin) for 12 h at 37°C. Tryptic digests were acidified (pH<3) using TFA and desalted cleaned up using iST cartridges (PreOmics, Munich) according to the manufacturer’s instructions. Samples were dried under vacuum and stored at −20 °C until further use.

### TMT labelling

Samples aliquots comprising 25 μg resp.12.5 μg of peptides were labelled with isobaric tandem mass tags (TMT 10-plex resp. TMTpro 16-plex, *Thermo Fisher Scientific*). The peptides were worked up and labelled as previously described^9^, although only half of the reported reagent volumes was used for the samples containing 12.5 μg peptides. Prior to the labelling reaction each sample was supplemented with a peptide calibration mixture consisting of six digested standard proteins mixed in different amounts, in order to control for ratio distortion during quantification. After stopping the labelling reaction with 2 M HCl the samples were further acidified using 5 % TFA, desalted using Sep-Pak Vac 1cc (50 mg) C18 cartridges (*Waters*) according to the manufacturer’s instructions and dried under vacuum.

### TMT LC-MS/MS analysis

TMT-labeled peptides were fractionated by high-pH reversed phase separation using a XBridge Peptide BEH C18 column (3,5 μm, 130 Å, 1 mm x 150 mm, *Waters*) on an *Agilent 1260 Infinity* HPLC system. samples were resolved with a flowrate of 42 μL/min using buffer A (20 mM ammonium formate in H_2_O, pH 10) and buffer B (20 mM ammonium formate in MeCN / 10% H_2_O (v/v), pH 10) and a linear gradient from 2%-10% (5 min), 10%-50% (55 min) (B) while monitoring peptide elution by UV absorption (215 nm, 254 nm). A total of 36 fractions were collected and pooled into 12 fractions using a post-concatenation strategy as previously described^10^ and dried under vacuum.

Dried peptides were resuspended in 0.1% aq. formic acid (v/v) and subjected to LC–MS/MS analysis using a Q Exactive HF Mass Spectrometer fitted with an EASY-nLC 1000 (both *Thermo Fisher Scientific*) and a custom-made column heater set to 60°C. Peptides were resolved using a RP-HPLC column (75 μm × 30 cm) packed in-house with C18 resin (*ReproSil-Pur* C18–AQ, 1.9 μm resin; *Dr. Maisch GmbH*) at a flow rate of 0.2 μL/min using Buffer A (0.1% Formic acid (v/v) in H_2_O) and Buffer B (0.1% Formic acid (v/v) in MeCN/20% H_2_O (v/v)) and with the following gradients: 5%-15% (10 min), 15%-30% (60 min), 30%-45% (20 min), 45%-95% (2 min) and 95% (18 min) (B) – for sample series labelled with TMT 10-plex 5%-15% (19 min), 15%-30% (80 min), 30%-45% (21 min), 45%-95% (2 min) and 95% (18 min) (B) – for sample series labelled with TMTpro 16-plex

The mass spectrometer was operated in DDA mode with a total cycle time of approximately 1 s. Each MS1 scan was followed by high-collision-dissociation (HCD) of the 10 most abundant precursor ions with dynamic exclusion set to 30 seconds. For MS1, 3e6 ions were accumulated in the Orbitrap over a maximum time of 100 ms and scanned at a resolution of 120,000 FWHM (at 200 m/z). MS2 scans were acquired at a target setting of 1e5 ions, maximum accumulation time of 100 ms and a resolution of 30,000 FWHM (at 200 m/z). Singly charged ions and ions with unassigned charge state were excluded from triggering MS2 events. The normalized collision energy was set to 30%, the mass isolation window was set to 1.1 m/z and one microscan was acquired for each spectrum.

### TMT LC-MS/MS data analysis

The acquired raw-files were analysed using the SpectroMine software (*Biognosis AG, Schlieren, Switzerland*). Spectra were searched against a murine database consisting of 17013 protein sequences (downloaded from Uniprot on 20190307) and 392 commonly observed contaminants. Standard Pulsar search settings for TMT 16 pro (“TMTpro_Quantification”) were used and resulting identifications and corresponding quantitative values were exported on the PSM level using the “Export Report” function. Acquired reporter ion intensities were employed for automated quantification and statistical analysis using our in-house developed SafeQuant R script v2.3^9^. This analysis included adjustment of reporter ion intensities, global data normalization by equalizing the total reporter ion intensity across all channels, summation of reporter ion intensities per protein and channel, calculation of protein abundance ratios and testing for differential abundance using empirical Bayes moderated t-statistics. Finally, the calculated p-values were corrected for multiple testing using the Benjamini–Hochberg method.

### PRM LC-MS/MS assay generation

In a first step, parallel reaction-monitoring (PRM) assays^11^ were generated from a mixture containing 50 fmol of each proteotypic heavy reference peptide of the target protein RSRP1 (AATEEASSR, ALGTTNIDLPASLR, MELLEIAK, TVYPEEHSR; JPT Peptide Technologies GmbH). Peptides were subjected to LC–MS/MS analysis using a Q Exactive Plus Mass Spectrometer fitted with an EASY-nLC 1000 (both Thermo Fisher Scientific) and a custom-made column heater set to 60°C. Peptides were resolved using a RP-HPLC column (75μm × 30cm) packed in-house with C18 resin (ReproSil-Pur C18–AQ, 1.9 μm resin; *Dr. Maisch GmbH*) at a flow rate of 0.2 μL/min using Buffer A (0.1% Formic acid (v/v) in H_2_O) and Buffer B (0.1% Formic acid (v/v) in MeCN/20% H_2_O (v/v)) and a linear gradient 5%-45% (60 min) (B).

The mass spectrometer was operated in DDA mode with a total cycle time of approximately 1 s. Each MS1 scan was followed by high-collision-dissociation (HCD) of the 20 most abundant precursor ions with dynamic exclusion set to 5 seconds. For MS1, 3e6 ions were accumulated in the Orbitrap over a maximum time of 254 ms and scanned at a resolution of 70,000 FWHM (at 200 m/z). MS2 scans were acquired at a target setting of 1e5 ions, maximum accumulation time of 110 ms and a resolution of 35,000 FWHM (at 200 m/z). Singly charged ions, ions with charge state ≥ 6 and ions with unassigned charge state were excluded from triggering MS2 events. The normalized collision energy was set to 27%, the mass isolation window was set to 1.4 m/z and one microscan was acquired for each spectrum.

The acquired raw-files were searched using the MaxQuant software (Version 1.6.2.3) against a human database (consisting of 20428 protein sequences downloaded from Uniprot on 20190307) using default parameters except protein, peptide and site FDR were set to 1 and Lys8 and Arg10 were added as variable modifications. The best 6 transitions for each peptide were selected automatically using an in-house software tool and imported into SpectroDive (version 8, *Biognosys, Schlieren*). A scheduled (window width 12 min) mass isolation lists containing the RSRP1 peptides was exported form SpectroDive and imported into the Q Exactive plus operating software for PRM analysis.

### PRM LC-MS/MS analysis

Peptide samples for PRM analysis were resuspended in 0.1% aqueous formic acid, spiked with the heavy reference peptide mix at a concentration of 4 fmol of heavy reference peptides per 1 μg of total endogenous peptide mass and subjected to LC-MS/MS analysis on the same LC-MS system described above using the following settings: The resolution of the orbitrap was set to 140,000 FWHM (at 200 m/z), the fill time was set to 500 ms to reach an AGC target of 3e6, the normalized collision energy was set to 27%, ion isolation window was set to 0.4 m/z and the first mass was fixed to 100 m/z. A MS1 scan at 35,000 resolution (at 200 m/z), AGC target 3e6 and fill time of 50 ms was included in each MS cycle. All raw-files were imported into SpectroDive for protein / peptide quantification. To control for variation in injected sample amounts, the total ion chromatogram (only comprising ions with two to five charges) of each sample was determined and used for normalization. To this end, the generated raw files were imported into the Progenesis QI software (*Nonlinear Dynamics* (*Waters*), Version 2.0), the intensity of all precursor ions with a charge of 2+ −5+ was extracted, summed for each sample and used for normalization. Normalized ratios were transformed from the linear to the log-scale, normalized relative to the control condition and the median ratio among peptides corresponding to one protein was reported.

### Code

The python scrpt used to map the eClip scores onto the proteomics data has been deposited to GitHub (https://github.com/Seedraking/eCLIP-Proteomics).

### qRT-PCR Measurement of Gene Expression

RNA was extracted from the treated cells by using TRI Reagent^®^ (*MRC*, TR118) according to the manufacturer’s protocol and reverse transcribed using the standard protocol for SuperScript™ III Reverse Transcriptase (*Thermo Fisher*), whereby Oligo(dT)15 Primers (*Promega*) were used. Measurement of the gene expression was done using PowerUP SYBR Green Master Mix (*Therno Fisher*) in triplicate (*n* = 3 biological triplicate) on an Applied Biosystems StepOnePlus Instrument with GAPDH as the housekeeping gene. The data was quantified using StepOne Software v2.3 and the −ΔΔ C_t_ values were used for averaging across the biological replicates.

### Primers used for qRT_PCR (5’-3’)

GAPDH_F CCACTCCTCCACCTTTGAC
GAPDH_R ACCCTGTTGCTGTAGCCA
KIF20A-a_F GCCCTGAAGAGCAATGAACGGGG
KIF20A-a_R ACCAGTACCCTCCATACCTGGGA
KIF20A-b_F AGCCGAGGGATCCTGACACCAC
KIF20A-b_R AGGCGGCATTTCTGAACGCGAA
KIF20B-a_F TTGAGTCTGAGGGCTGTGTGCA
KIF20B-a_R TCTGTGCCATCTGCCCTGAGCT
KIF20B-b_F TCCTTATGCTTCCCACCTGTGA
KIF20B-b_R AGAACTACTCTGGGGAGTTCCA
RSRP1-a_F GCTACTCGCGGTCATACTCGC
RSRP1-a_R GGTACCGCGAAGGAGACCGGTA
RSRP1-b_F CTTTCCGGAGAAAGCGCAGGC
RSRP1-b_R CGAATCCTTCTCCTGCGGCGAG
RSRP1-c_F TGTGTAATTGGGCCCAGATGTGG
RSRP1-c_R CCAGTTAGGCACCTAATCGTCTGA
RSRP1-d_F CCTACCCAGCAAAGAAGCATAGCT
RSRP1-d_R TGCCTCTTCTGTGGCAGCTTTAGC
RSRP1-e_F AGCCAAAGAAACAAGCCGTGGA
RSRP1-e_R TGTCTGCAGACCACATTCATGA
RSRP1-f_F TGGCTAAGTTTGCATGAAAACTGCA
RSRP1-f_R CCCACCTGTGCATAACCTTTGGTCC
RSRP1-g_F AGGAAGTCCTGAGTTGAGG
RSRP1-g_R GTCGTTCACGTAGTTGGAC
RBM39-b_F GCCAGCAGCACAGCAAGCTCTA
RBM39-b_R GCTGAACAGAGGCAGCTGCAGC
RBM23_F CAGTTAGCTGCCCGAATTCGGCC
RBM23_R ACGTACATCGCGAACCTTGCCT

### CRISPR-Cas9 mediated HCT-116 DCAF15−/− cells

The cells were generated using previously reported strategies. Briefly, HCT-116 cells in 2 wells (6-well plate) were transfected with the plasmids and first selected with puromycin for plasmid transfection and then with indisulam for CRISPR-Cas9 induced DCAF15 editing. Hereafter, the cells were diluted and re-seeded in multiple 96-well plates such that there would be one cell per 20 wells on average (visual inspection showed higher dilution). The cells were grown in DMEM with 10% FCS, 1% penicillin/ streptomycin and 3 uM indisulam. Colonies from 2 wells were picked and expanded seperately. PCR analysis and Sanger sequencing confirmed homozygous knock out of the targeted gene fragment for both isolated clones. Western Blotting confirmed lack of RBM39 degradation upon indisulam treatment. The growing medium was regularly changed across all the stages.

### siRNA

The siRNA-mediated knockdowns of DCAF15 (J-031237-18, *Horizon Discovery*), RBM23 (J-016689-11, *Horizon Discovery*) and RBM39 (D-011965-02, *Horizon Discovery*) in HCT-116 cells were done using the reverse transfection protocol for RNAiMax (*Thermo Fisher*), whereby a solution of siRNA (3 μL, 10 μM) diluted in 197 μL Opti-MEM I (*Gibco*) was directly added to 2 μL of RNAiMAX in the well. The mixture was left to incubate at rt for 5 min after which a suspension of 200 k cells in 0.8 mL DMEM supplemented with 10% FCS was added. The cells were left to incubate at 37 °C with 5% CO_2_ for 72h or 96h.

### Flow cytometry

500k HCT-116 cells were were seeded in a 6 well-plate format in 2 mL growing medium. For cell synchroization a double thymidine block was conducted by treating the cells the next day with 40 μL thymidine (100 mM in PBS) and left to grow in the incubator for 16 h. Hereafter, the growing medium was removed and the cells washed with warm PBS once. The cells were left too grow in fresh medium for another 8 h and then treated with 40 μL thymidine again. Once they had grown for another 16 h, the medium was removed again, the cells washed with warm PBS. Subsequently, they were grown (released) and treated in fresh medium for the indicated time points. The growing medium was removed and 0.05% Trypsin was added to detach the cells. The trypsin was neutralizied by the addition of growing medium and the cells collected via centrifugation (200 *g* at 4°C for 5 min). The pellets were washed twice with PBS, resuspended in cold PBS and fixed with 70% ethanol. The fixed cells were collected by centrifugation (8000k *g* at 4°C for 5 min), resuspended in staining buffer (0.1 % (v/v) Triton; 0.005% Propidium Iodide; 0.02% DNAse-free RNAse A). The cells were incubated at 37 °C for 15 min after which they were measured on a BD LSR Fortessa Analyzer and the data analyzed with FlowJo software (version 10.5.3, TreeStar).

### Immunofluorescence

HCT-116 cells were seeded on coverslips in a 6 well-plate or 12 well-plate format in 2 mL resp. 1 mL growing medium. They were treated with DMSO, E7820 or GSK-923295 at the indicated concentrations for the indicated time. The cells were washed twice with warm PBS, fixed using 3.7% Paraformaldehyde in PBS and were permeabilized with 0.1% (v/v) Triton in PBS. The coverslips were blocked in 10% BSA in PBS at rt for 1h and then incubated with antibodies for CENPE-E (Abcam; ab5093, 1:500 dilution) and α-Tubulin (GeneTex, GTX112141, 1:250 or Abcam, ab7291, 1:500 dilution) at rt for 1h. The coverslips were further developed using anti-Mouse (Thermo Scientific, Alexa Fluor 594, 1:200 dilution) and anti-Rabbit secondary antibody (Thermo Scientific, Alexa Fluor 488, 1:200 dilution) at rt for 1h. Finally, they were incubated with DAPI (0.2 μg/ml) for 5 min and mounted on slides using ProLong™ Diamond Antifade Mountant. The images were collected at 0.2 μm z-sections with a ×60 1.35 NA oil objective using a DeltaVision Core system (Applied Precision) with a Coolsnap HQ^2^ camera (Roper). Images were deconvolved with SoftWoRx (Applied Precision) and maximum-intensity two-dimensional projections were assembled using FIJI (ImageJ, NIH). For the quantification, 100 cells were analyzed for each treatment condition and classified into mitotic, non-mitotic and multi-nucleated cells. Each quantification was conducted in a biological triplicate (*n=3*).

### Chemical Synthesis

#### General

Unless otherwise stated, all chemicals and solvents were purchased from commercial sources and used as received. Reactions were monitored by analyzing a small sample (after suitable workup) by TLC or UHPLC-MS. For chromatographic purifications HPLC-grade solvents and nanopure water were used. Evaporation of the solvents and volatiles *in vacuo* was done with the rotary evaporator. Aqueous solutions were frozen and lyophilized.

#### Chromatograpic purification

Flash chromatography was carried out on a *Biotage* Isolera Four with self-packed columns. For reverse phase *LiChroprep RP-18*, 40-63 μM silica (Merck) was used with the indicated solvent system, whereby the gradients were adjusted for the individual compounds based on their UHPLC trace.

Preparative RP-HPLC was performed on a Shimadzu Prominence UFLC Preparative Liquid Chromatograph using a *Gemini NX-C18*, 5 μm, 110 Å, 21.2 x 250 mm from Phenomenex column with a flow rate of 20 mL/min and the indicated solvent system, whereby the gradients were adjusted for the individual compounds based on their UHPLC trace.

Solvent systems used for RP-flash chromatography and preparative RP-HPLC: System A) Buffer A (0.1% TFA (v/v) in H_2_O), Buffer B (0.1% TFA (v/v) in MeCN) System B) Buffer A (H_2_O), Buffer B (MeCN)

#### Analysis

##### TLC

*Merck* tlc plates silica gel *60* F254, 0.25 mm pre-coated glass plates; the spots were visualized by UV light (254 and 366 nm) or using staining reagents.

##### UHPLC-ESIMS

*Agilent 1290 Infinity* system equipped with a C_18_ column (*ZORBAX Eclipse Plus RRHD*, 1.8 μm) and attached to an *Agilent 6130 Quadrupole*; samples were run with a flowrate of 0.45 mL/min at 40°C with Buffer (A): 0.1% formic acid (v/v) in H_2_O/1% MeCN (v/v), Buffer (B): 0.1% formic acid (v/v) in MeCN/1% H_2_O (v/v) and a gradient 5-90% (3.5 min), 90% (1 min) (B).

##### NMR

^1^H, ^13^C and 2D-NMR were recorded on a *BrukerAvance* (400 or 500 MHz); δ in ppm relative to solvent signal; multiplicities in Hz (reported as br = broad signal, s = singlet, d = doublet, t = triplet or a combination of these eg. dd).

##### HRMS

*Bruker maXis 4G QTOF* ESI mass spectrometer.

#### Synthesis of SC_046

As previously reported[8] SC_046 was synthesized by dissolving 7-nitro indole (1.0 g, 6.17 mmol) in THF (10 mL) followed by the addition of aq. HCl (20 μL, 0.5 M) and N-chlorosuccinimide (840 mg, 6.3 mmol). The reaction mixture was stirred at rt overnight after which H_2_O was added. The crystalline precipitate was filtered, washed with H_2_O, MeOH-H_2_O (1:1), isopropylether and dried *in vacuo* to yield SC_046 (1.22 g, quantitative).

*Data* for **SC_046 - ^1^H-NMR** (CDCl_3_, 400 MHz): δ 9.72 (br, 1H), 8.07-8.12 (d, *J* = 8.0 Hz, 1H), 7.84-7.91 (d, *J* = 7.9 Hz, 1H), 7.15-7.19 (t, *J* = 7.9 Hz, 1H), 7.13 (s, 1H).

#### Synthesis of SC_047

As previously reported[8] SC_047 was synthesized by dissolving SC_046 (500 mg, 2.5 mmol) in 2-propanol (7 mL) and adding Fe powder (426 mg, 7.6 mmol) as well as a solution of NH_4_Cl (27.2 mg, 0.5 mmol) in H_2_O (1.3 mL) to it. The reaction mixture was stirred at 60 °C for 4 h after which activated charcoal (200 mg) was added. The insolubles were filtered over celite and washed with EtOAc (100 mL). The filtrate was reduced *in vacuo* yielding SC_047 (424 mg, 89%) which was used without further purification.

*Data* for **SC_047 - ^1^H-NMR** (CDCl_3_, 400 MHz): δ 7.15-7.19 (d, *J* = 8.1 Hz, 1H), 7.13-7.15 (d, *J* = 2.6 Hz, 1H), 7.01 −7.06 (t, *J* = 7.5 Hz, 1H), 6.62-6.66 (dd, *J* = 7.4 Hz, *J* = 0.8 Hz, 1H).

#### Indisulam

Indisulam was synthesized from SC_047 as previously described[64] using 4-sulfamoylbenzene-1-sulfonyl chloride and purified by preparative RP-HPLC (Solvent system A) yielding the product (120 mg, 80%).

*Data* for **Indisulam - ^1^H-NMR** (CD_3_OD, 500 MHz): δ 7.97-7.95 (m, 2H), 7.82-7.81 (m, 2H), 7.38-7.36 (dd, *J* = 1.0, 9.0 Hz, 1H), 7.27 (s, 1H), 6.92-6.89 (t, *J* = 7.7 Hz, 1H), 6.61 −6.59 (dd, *J* = 0.8, 7.6 Hz, 1H). ^13^**C-NMR** (CD_3_OD, 125 MHz): δ 149.1, 143.8, 132.4, 129.1, 128.5, 127.8, 123.5, 122.3, 120.9, 119.8, 106.3. **UHPLC-ESIMS** (m/z): 374 [M]-H^+^.

#### Synthesis of E7820

To a solution of 7-amino-4-methyl-1*H*-indole-3-carbonitrile (15 mg, 0.083 mmol) in THF (1 mL) was added 3-Cyanobenzenesulfonyl chloride (23 mg, 0.115 mmol) and pyridine (21 μL) under inert atmosphere. The mixture was stirred at rt for 3 h after which the volatiles were removed *in vacuo* and the crude purified with preparative RP-HPLC (System A) to yield E7820 (27.4 mg, 98%).

*Data* for **E7820 - ^1^H-NMR** (DMSO-d_6_, 500 MHz): δ 12.02 (d, *J* = 2.3 Hz, 1H), 10.13 (s, 1H), 8.18 (d, *J* = 3.2 Hz, 1H), 8.13–8.08 (m, 2H), 7.95–7.89 (m, 1H), 7.76–7.70 (m, 1H), 6.80 (dd, *J* = 7.7, 0.8 Hz, 1H), 6.53 (d, *J* = 7.7 Hz, 1H), 2.58 (s, 3H. **^13^C-NMR** (DMSO-d_6_, 125 MHz): δ 140.4, 136.5, 135.5, 131.4, 131.3, 130.6, 130.5, 128.4, 126.6, 122.5, 119.6, 119.6, 117.4, 117.3, 112.4, 84.4, 17.7. **UHPLC-ESIMS** (m/z): 337 [M]+H^+^.

#### Chloroquinoxaline sulfonamide (CQS)[13]

CQS was synthesized 2,5-dichloroquinoxaline and 4-aminobenzene-1-sulfonamide as previously reported^4^ and purified by flash chromatography (pentane: ethylacetate) to yield CQS (55mg, 33%).

Data for CQS - **^1^H-NMR** (DMSO-d_6_, 500 MHz): δ 11.70 (br, 1H), 8.62 (s, 1H), 7.78-7.67 (m, 5H), 6.60-6.57 (m, 2H), 6.07 (s, 2H), 5.74 (s, 1H). **UHPLC-ESIMS** (m/z): 335 [M]+H^+^.

#### Tasisulam[65]

Tasisulam was synthesized from 5-Bromothiophene-2-sulfonamide and 2,4-Dichlorobenzoyl chloride as previously reported^5^ and purified by preparative RP-HPLC (System B) to yield Tasisulam (82mg, 95%).

Data for Tasisulam - **^1^H-NMR** (CD_3_CN, 500 MHz): δ 10.12 (br, 1H), 7.68-7.67 (d, *J* = 4.1 Hz, 1H), 7.54-7.53 (d, *J* = 2.0 Hz, 1H), 7.48-7.46 (d, *J* = 8.3 Hz, 1H), 7.41 −7.39 (dd, *J* = 2.08, 8.3 Hz, 1H), 7.25 (d, *J* = 4.1 Hz, 1H). **^13^C-NMR** (CD_3_CN, 125 MHz): δ 164.7, 140.4, 138.4, 136.5, 133.0, 132.5, 132.1, 131.5, 130.9, 128.5, 122.9. **UHPLC-ESIMS** (m/z): 414 [M]+H^+^.

